# On-Pathway Oligomer of Human Islet Amyloid Polypeptide Induced and Stabilized by Mechanical Rotation During MAS NMR

**DOI:** 10.1101/2023.07.06.547982

**Authors:** Samuel D. McCalpin, Malitha C. Dickwella Widanage, Riqiang Fu, Ayyalusamy Ramamoorthy

**Author notes:** Corresponding Author Correspondence and requests for materials should be addressed to A.R.

## Abstract

Intermediates along the fibrillation pathway are generally considered to be the toxic species responsible for the pathologies of amyloid diseases. However, structural studies of these species have been hampered by heterogeneity and poor stability in standard aqueous conditions. Here, we report a novel methodology for producing stable, on-pathway oligomers of the human Type-2 Diabetes-associated islet amyloid polypeptide (*h*IAPP, or amylin) using the mechanical forces associated with magic angle spinning (MAS). The species were a heterogeneous mixture of globular and short rod-like species with significant β-sheet content and the capability of seeding *h*IAPP fibrillation. We used MAS NMR to demonstrate that the nature of the species was sensitive to sample conditions including peptide concentration, ionic strength, and buffer. The methodology should be suitable for studies of other aggregating systems.

---

Several degenerative diseases, including Alzheimer’s Disease (AD), Parkinson’s Disease (PD), and Type-2 Diabetes (T2D), are linked to the aggregation of proteins and peptides into fibrillar structures called amyloid.^1–4^ While the relationship between amyloid aggregation and these diseases was established several decades ago, few successful treatments have been developed that target the fibril formation process.5 Structural biologists seeking to guide drug design struggle to characterize amyloid fibrils because their large size and non-crystalline nature render traditional solution NMR and crystallography unfeasible. Still, solid-state NMR and Cryo-EM have proven capable of revealing specific, atom-resolution cross-β-sheet structures in fibril cores. However, recent research indicates that intermediate aggregates along the amyloid formation pathway, called “oligomers” or “protofibrils” based on size and morphology, are the toxic species most responsible for the pathologies of AD, PD, T2D, and other amyloidoses.^6–9^ Structural characterizations of oligomeric amyloid aggregates would greatly benefit efforts to develop treatments against amyloid diseases, but several factors have made them scarce if not nonexistent. The most significant of these factors are high heterogeneity among oligomeric species and their short lifetimes in aqueous conditions.

Magic angle spinning (MAS) solid-state NMR spectroscopy is uniquely positioned to characterize heterogeneous systems of intermediate molecular weight, slow-tumbling species like amyloid oligomers. Its sensitivity to individual nuclei in distinct chemical environments allows discrimination between differently structured conformers. Additionally, a combination of a dipolar recoupling experiment and intermediate frequency MAS has been demonstrated to facilitate the separation of the oligomer signal from that of monomers and fibrils in a heterogeneous preparation of amyloid-beta.^10^ Typically, MAS experiments require biological samples to be lyophilized, frozen, or crystallized, but comprehensive multiphase (CMP) NMR spectroscopy enables the direct observation of mixed-phase systems with liquids.^11^ The technique has been used to study microorganisms and environmental samples but also presents an approach to characterize heterogeneous amyloid systems consisting of soluble fast-tumbling monomers and low-order oligomeric aggregates and insoluble slow-tumbling high-order oligomers and fibrillar aggregates. The significant mechanical forces involved in MAS additionally offer a novel modulator of the amyloid formation pathway. Previous work has demonstrated the ability of small molecules, peptides, lipid membranes, metals, synthetic polymers, pH, and electric fields to alter amyloid aggregation pathways and in some cases to stabilize oligomeric intermediates for further structural studies.^12–24^ Here, we provide the first report, to our knowledge, of the ability of the mechanical forces associated with MAS to stabilize intermediate, on-pathway aggregates of human islet amyloid polypeptide (*h*IAPP) in an aqueous environment.

To compare the aggregation behavior of *h*IAPP with and without MAS, we prepared two samples and collected ^1^H NMR spectra over the course of 24 hours using a standard static solution probe and a CMP probe with 10 kHz MAS (**Figure 1**). The sample conditions were identical except for 100% D_2_O used in the MAS sample compared to 10% D_2_O used in the solution NMR sample. Under the conditions of the solution NMR experiment, the ^1^H NMR spectrum of *h*IAPP appeared with the lineshape in **Figure 1a**, and the signal quickly decayed as the peptide aggregated within 7-8 hours. In contrast, the MAS sample immediately afforded a much broader lineshape with a significant peak near 0 ppm, typical of aggregated *h*IAPP (**Figure 1b**). This signal was also stable for at least 24 hours (**Figure 1c**). To rule out D_2_O as the cause for the differences between the samples, we performed a thioflavin T fluorescence assay with buffers of varying amounts of D2O and observed no change in the kinetics based on the percentage of D2O (**Figure S1**). Thus, our data suggests that MAS induced the formation of an *h*IAPP aggregate.

**Figure 1.**
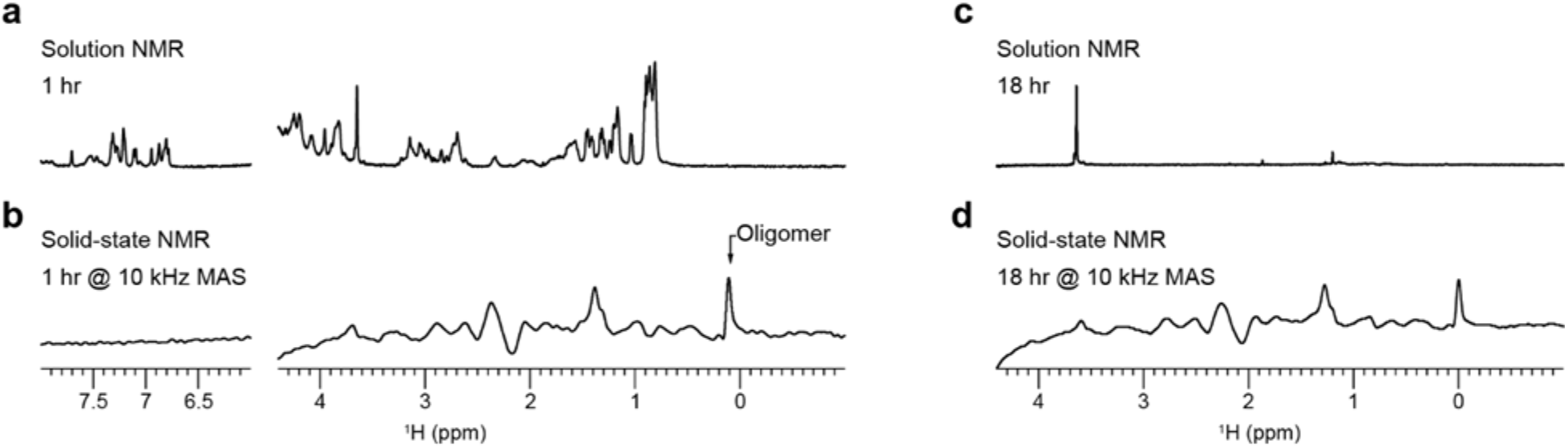
Visualizing the oligomeric intermediate’s peaks under MAS. *h*IAPP monomers were detected from solution NMR (**a, c**) and MAS NMR (**b, d**) using 1H spectra obtained at 1- and 18-hour timepoints on a 500 MHz spectrometer at 298 K. Samples contained 50 mM *h*IAPP in 10 mM d11-tris, 100 mM NaCl, pH 7.4 buffer.

However, due to the smaller volume of the MAS rotor, significantly less material was present in the MAS experiment than in the solution experiment, causing a noisier signal. There was also notable baseline rolling in the MAS spectrum which we hypothesized arose from conductivity through the coil due to salt in the buffer. To address these issues, we performed the MAS experiment again using a sample with a higher concentration of *h*IAPP in pure D_2_O (**Figure 2a**). Interestingly, the aggregation behavior of this sample, as assessed by a series of ^1^H NMR spectra collected under 10 kHz MAS, differed greatly from that of the previous sample. In this case, the ^1^H NMR lineshape was initially much sharper and decayed over time to a broader, less intense lineshape which was stable after 18 hours. Subtracting the spectrum at 18 hours from the initial one revealed a lineshape which was remarkably similar to that of the solution spectrum, suggesting that the spectral changes over time resulted from the depletion of *h*IAPP monomers as they associated into larger species. Additionally, we subtracted the solution spectrum from the long-time MAS spectrum, normalized to the intensity of the peak at 0.95 ppm, to obtain the lineshape of the *h*IAPP aggregates. The presence of a peak near 0 ppm, which did not vary with time, suggested that these species (denoted as MAS-*h*IAPP) were formed immediately under MAS and were stable for at least 24 hours.

**Figure 2.**
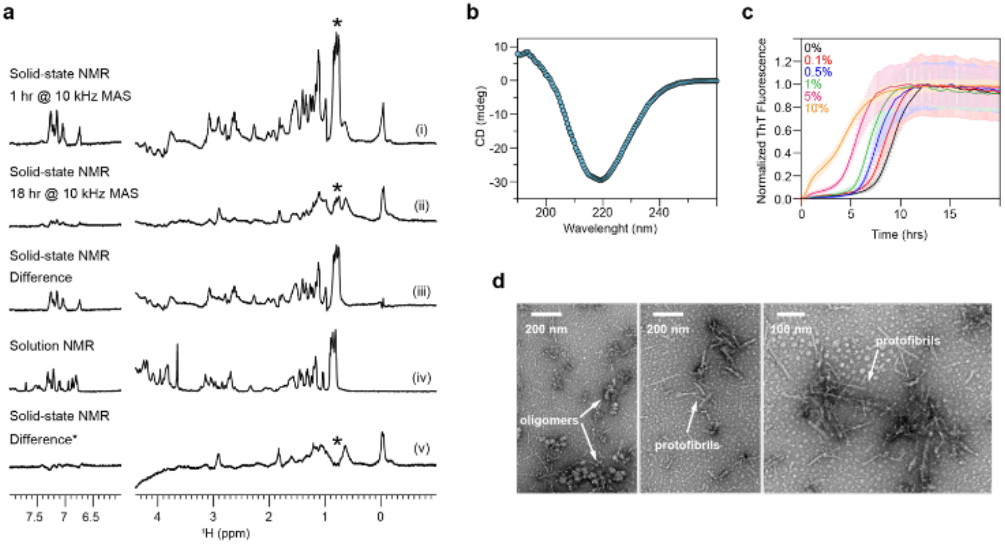
Heterogeneous, b-sheet structured, on-pathway *h*IAPP oligomers induced by MAS. **(a)** 1D ^I^H NMR spectra with spectral editing showing the MAS-hIAPP oligomeric intermediates: **(i)** after 1 hour, **(ii)** after 18 hours, **(iii)** the difference spectrum of i and ii (normalized by the number of scans), **(iv)** solution NMR spectrum of *h*IAPP, and **(v)** the difference spectrum of ii and i (normalized by the peak with asterisk). **(b)** b-sheet structure identified using CD spectroscopy. **(c)** ThT fluorescence assay of *h*IAPP monomers with noted concentrations of MAS-*h*IAPP seeds. **(d)** TEM revealed heterogeneous MAS-*h*IAPP species; oligomers and protofibrils are labeled. For MAS NMR, 80 nmol *h*IAPP was hydrated to 2 mM in D_2_O. NMR data were collected over 48 hours with 10 kHz MAS, and then the sample was removed from the rotor for further characterization. The MAS-*h*IAPP was diluted to 80 mM in D2O for CD spectroscopy, 100 mM for TEM, and the noted molar ratios relative to 5 mM monomeric *h*IAPP in sodium phosphate buffer (10 mM NaPO4, 100 mM NaCl, 10 mM ThT, pH 7.4) for ThT experiments.

We then characterized the MAS-*h*IAPP by a combination of fluorescence, CD spectroscopy, and TEM. CD exhibited a strong negative peak at 218 nm, indicative of significant bsheet content (**Figure 2b**). The ThT fluorescence assay showed sigmoidal aggregation kinetics for *h*IAPP monomers in the presence of seeds taken from the MAS sample. In contrast, the fluorescence intensity of ThT in the presence of the MAS aggregate alone was constant over time and noticeably less than that of the monomers which formed fibrils (**Figure 2c**). Lag times of monomer aggregation decreased with increasing concentrations of MAS-*h*IAPP seeds, demonstrating that the MAS-*h*IAPP seeded aggregation of *h*IAPP monomers. Taken together, these results indicate that MAS induced the formation of on-pathway *h*IAPP oligomers with significant b-sheet content. However, TEM revealed a mixture of morphologies, including amorphous globular aggregates and rod-like protofibrillar species which were much shorter than the fibers formed by *h*IAPP under static conditions (**Figure 2d**). Based on the data presented here, we cannot determine whether one or both observed species are on-pathway. But it seems most likely that the smaller globular aggregates were the dominant contributors to the NMR and CD spectra, assuming their faster tumbling and greater aqueous solubility. Regardless, the data confirms that MAS induced the formation of non-fibrillar oligomeric intermediates of *h*IAPP.

There were issues reproducing the MAS-*h*IAPP formation in D_2_O (**Figure S2**), perhaps due to the use of an unbuffered solvent. Solution pH has been extensively reported to affect the kinetics of *h*IAPP aggregation and the end species formed, so it is favorable to control regardless of any issues with reproducibility.^25–28^ To this end, we prepared *h*IAPP samples in several buffer conditions and collected ^1^H MAS spectra as before. The observed ^1^H lineshapes for each sample condition were compiled in **Figure 3**. In all cases, there was a peak near 0 ppm at the initial timepoint which did not significantly change for at least 18 hours, consistent with the immediate formation of a stable oligomer regardless of buffer. The 0 ppm peak was weakest for the low *h*IAPP concentration, acetate buffer condition, which is unsurprising given that both low peptide concentration and low pH are known to reduce *h*IAPP aggregation (**Figure 3f**).^26,29^ Remarkably, though there was significant spectral variation across the samples, the lineshapes reported for the oligomers formed in 2 mM *h*IAPP in pure D_2_O, Tris buffer (no salt), and sodium phosphate buffer (with salt) were very similar (**Figure 3a-c**). This suggests that a similar species, or range of species, formed under these sample conditions. Based on the differences between the spectral patterns of **Figures 3d** and **3e** and between **Figures 3a, 3b**, and **3d**, the peptide concentration and buffer ionic strength seemed to be the most significant factors which influenced the nature of the oligomers formed. Tris also appeared to interact with *h*IAPP and bias the aggregation pathway, but only in the presence of salt.

**Figure 3.**
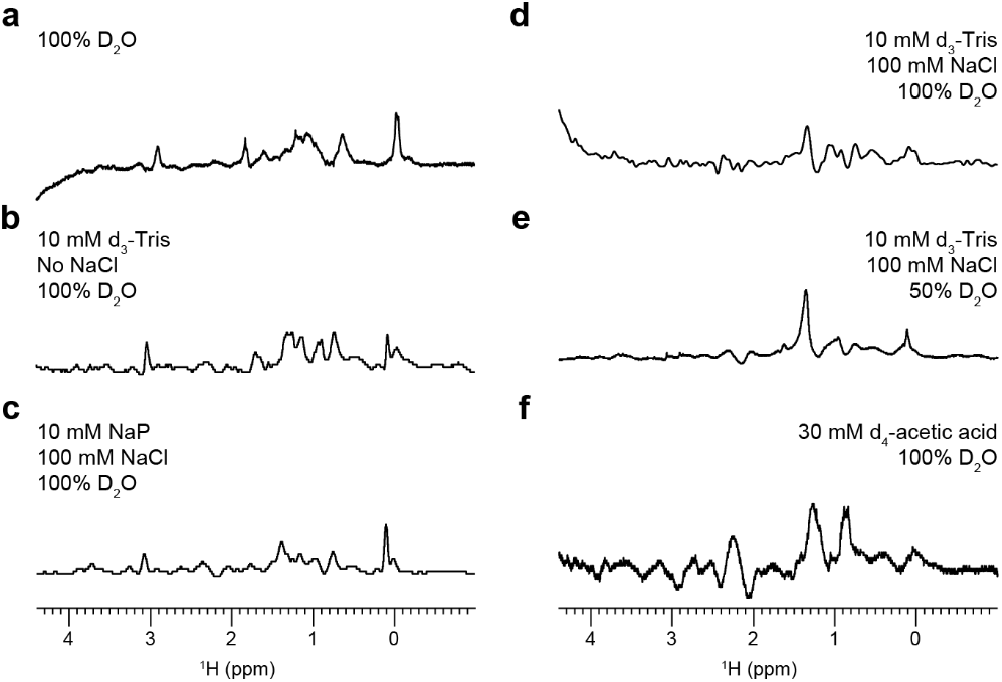
MAS-induced aggregation pathway altered by buffer. Solution conditions modified *h*IAPP aggregation under 10 kHz MAS. *h*IAPP concentrations are **(a)** 2 mM, **(b)** 2 mM (7.4 pH), **(c)** 2 mM (7.4 pH), **(d)** 2 mM (7.4 pH), **(e)** 50 μM (7.4 pH), and **(f)** 50 μM (5.5 pH). A 0-ppm peak indicated the presence of oligomeric intermediates.

The amount of structural information we could obtain from 1D ^1^H NMR lineshapes was limited, so we collected 2D ^1^H-^1^H RFDR MAS spectra of a MAS-*h*IAPP sample in pure D_2_O (**Figure 4**).^30^ These RFDR spectra (**Figure 4a**) exhibited several well-resolved cross-peaks which mostly saturated with 25 ms mixing (**Figures 4c, S3**). A comparable spectrum of MAS-*h*IAPP in buffer (**Figure S4**) displayed significantly fewer (but still well-resolved) cross-peaks. Considering that the primary difference between these two samples was a significant monomer population present in pure D_2_O and not in buffer, the extra cross-peaks in the spectrum of the D_2_O sample likely arose from this population. It seems unlikely that dipolar recoupling would occur so efficiently in small, fast-tumbling monomers, so it is also possible that the observed cross-peaks represented low-order oligomers or monomers which were motionally restricted by transient interactions with larger, invisible oligomers. Regardless, the RFDR spectra demonstrated that MAS-*h*IAPP samples should be amenable to structural studies. Challenges related to peak assignments could be overcome in the future with 3D heteronuclear experiments on ^13^C-labeled peptide, and higher MAS frequency could allow full observation of the larger oligomers.

**Figure 4.**
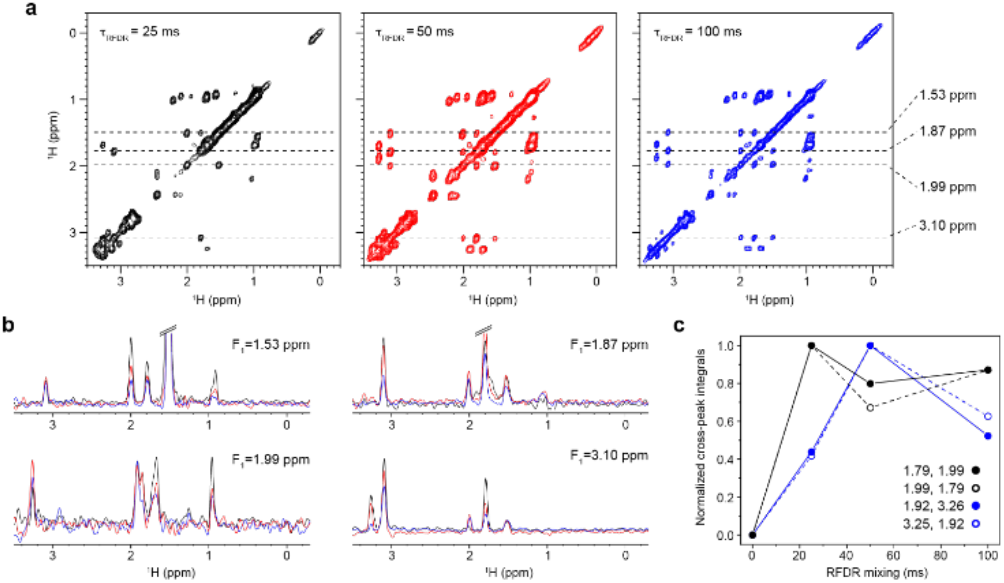
Dipolar recoupling from MAS-*h*IAPP. 2D ^1^H-^1^H RFDR spectra of 2 mM *h*IAPP in 100% D_2_O obtained with an 800 MHz NMR spectrometer under 15 kHz MAS at 298 K. **(a)** Spectra were obtained at three mixing times. **(b)** 1D slices and **(c)** build-up curves are plotted for the cross-peaks at the noted chemical shifts.

Drug design efforts against amyloid diseases would greatly benefit from structural studies of intermediate amyloid aggregates. Here, we presented a novel method to stabilize nonfibrillar species of *h*IAPP using the mechanical forces associated with MAS. We showed that the heterogeneous mixture of species included rod-like protofibrils and globular aggregates and on-pathway, beta-sheet-rich species which formed within several hours in pure D_2_O, or immediately in common buffer conditions. The sample conditions, namely the peptide concentration and choice of buffer, were also found to affect the oligomers formed. 2D RFDR experiments showed that the MAS-*h*IAPP was amenable to further structural characterization. Such studies would deepen our understanding of the molecular basis of T2D. However, their feasibility in our hands was limited by the availability of isotope-labeled peptide and MAS limits on liquid samples. Significantly, the methodology reported here should be broadly applicable to studying aggregates of other amyloidogenic peptides and IDPs and is more tunable than chemical chaperones of oligomeric aggregates.

## ASSOCIATED CONTENT

### Supporting Information

Methods and materials, additional ThT fluorescence and MAS NMR data, discussion of MAS NMR data reproducibility. This material is available free of charge via the Internet at http://pubs.acs.org.

## Author Contributions

The manuscript was written through contributions of all authors.

## ACKNOWLEDGMENT

This work was supported by the NIH grant 5R01DK132214-02. A portion of this work was performed at the National High Magnetic Field Laboratory, which is supported by National Science Foundation Cooperative Agreement No. DMR-2128556 and the State of Florida.

## ABBREVIATIONS

*h*IAPP: human islet amyloid polypeptide
NMR: nuclear magnetic resonance
MAS: magic angle spinning

## SYNOPSIS TOC

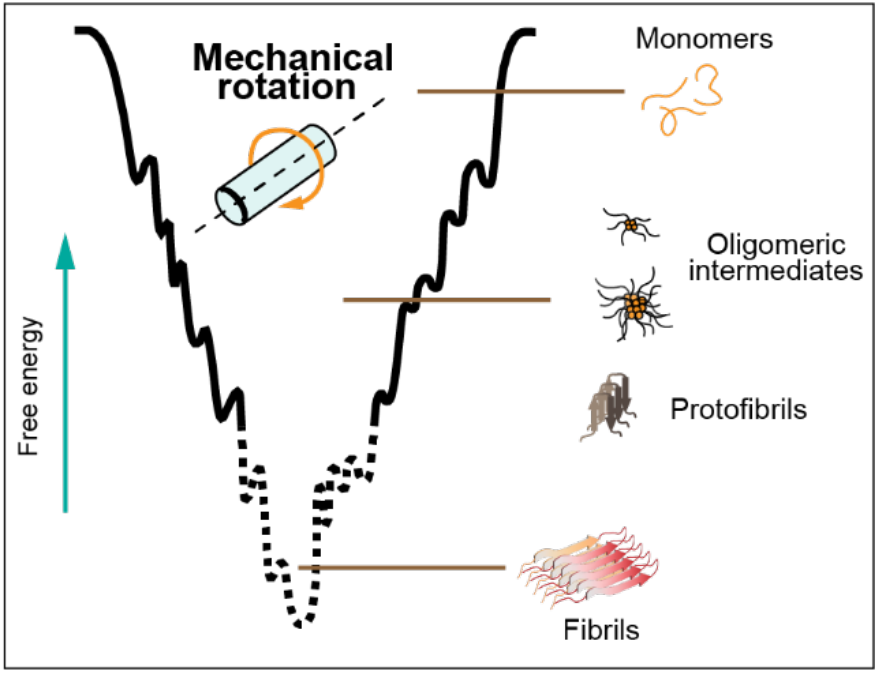

