## Supporting Information for "On-Pathway Oligomer of Human Islet Amyloid Polypeptide Induced and Stabilized by Mechanical Rotation During MAS NMR"

### Materials and Methods

*Materials*: Synthetic, amidated human IAPP was purchased from Anaspec, treated with hexafluoro-isopropanol for one hour at room temperature, lyophilized, and stored at -20 °C prior to use. All other chemicals were obtained from Sigma Aldrich.

*NMR Experiments*: For MAS NMR at 500 MHz, 80 nm human islet amyloid polypeptide (*h*IAPP) was dissolved in HFIP and lyophilized directly into a KF 4 mm MAS rotor insert. The peptide was hydrated in 40 µL of solvent, the rotor insert was inserted in a 4 mm rotor, and experiments were performed on Bruker 500 MHz (11.7 T) spectrometer using a 4 mm Comprehensive MultiPhase (CMP), triple-resonance HCN probe under 10 kHz MAS at 298 K. The radiofrequency field strengths were 55.5 kHz for $^{1}H$ hard pulse used for excitation. Sample conditions are specified under each figure caption.

Experiments at 800 MHz were conducted with a Bruker 800 MHz (18.8 T) and a 3.2 mm homemade HX MAS probe. Samples were prepared as above and 20 µL was transferred into a 3.2 mm rotor under 15 kHz MAS at 298 K.

*Solution NMR*: $^{1}H$ NMR spectra of *h*IAPP (50 µM in 10 mM $d_{11}$-tris, 100 mM NaCl, 10% $D_2O$ for locking, pH 7.4 buffer) were collected on a 500 MHz Bruker NMR spectrometer using a triple-resonance TX1 probe. Spectra are presented as an average of 1024 scans collected with a 13 µs 90° pulse and a 3 s recycle delay.

*TEM*: After 48 hours of spinning at 10 kHz, *h*IAPP (2 mM, 100% $D_2O$) was removed from the NMR rotor and diluted to a final concentration of 100 µM in $D_2O$. For TEM, 5 µL of the *h*IAPP sample was blotted on a carbon holey-mesh grid, and the grid was stained with 10 µL of 2% uranyl acetate. Images were collected on a JEOL TEM.

*CD Experiments*: The MAS NMR *h*IAPP sample in 100% $D_2O$ was diluted to 80 µM in $D_2O$ and its CD spectrum was measured in a JASCO CD-spectropolarimeter using an average of 10 accumulations with 1 nm bandwidth, 0.5 nm data pitch, 100 nm/minute scanning speed, 1 s data integration time, and 200 mdeg CD scale.

*Seeding Experiments*: The *h*IAPP in 100% $D_2O$ sample was collected after solid-state NMR experiments (48 hours of 10 kHz MAS) and diluted in $D_2O$. These MAS *h*IAPP species were mixed in several molar ratios with freshly prepared monomeric *h*IAPP (5 µM) and thioflavin-T (10 µM) in sodium phosphate buffer (10 mM $NaPO_4$, 100 mM NaCl, pH 7.4). Samples (50 µL) were plated in quadruplicate in a black-walled, flat-bottomed 384-well microplate (Gruyner) and fluorescence emission at 480 nm was after exciting at 454 nm. The ThT fluorescence experiments were performed at 25° C with measurements collected every 8 minutes.

**Supplemental Figures**

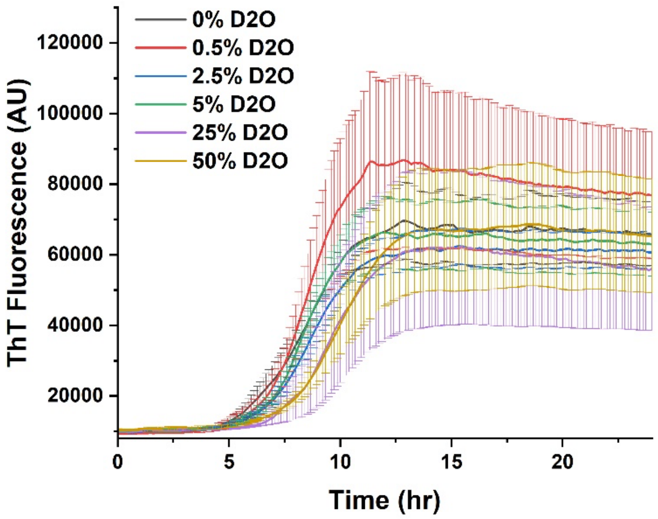


**Figure S1. $D_2O$ does not affect *h*IAPP aggregation.** ThT fluorescence measured over time for 5 µM *h*IAPP in tris buffer (10 mM Tris, 10 µM ThT, 100 mM NaCl, pH 7.4) with the noted $D_2O$ compositions.

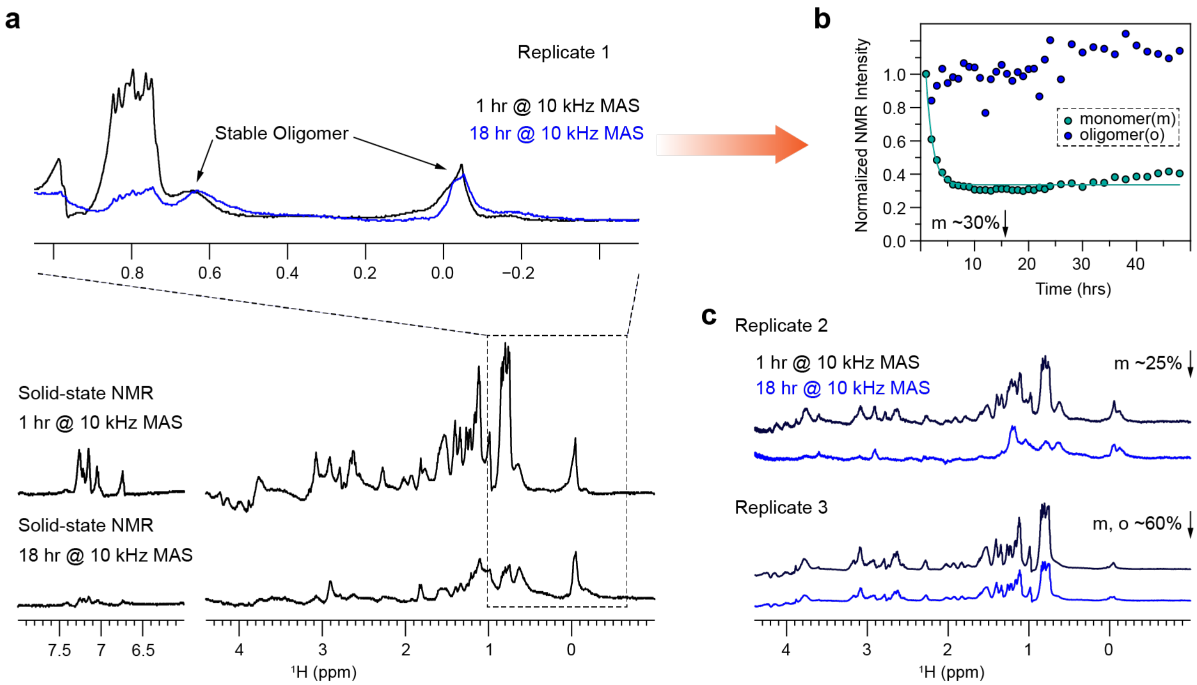


**Figure S2. Oligomeric intermediates show stability, but reproducibility is challenging. a,** MAS-induced oligomeric intermediates are stable after 18 hours under 10 kHz MAS. 1 ppm to -0.4 ppm region is zoomed in. **b,** Normalized intensities of 1 ppm to 0.5 ppm (represent monomers) and 0.2 ppm to -0.3 ppm (represent oligomeric intermediates) in 1D $^{1}$H spectra show monomers are depleted up to 30% while formed oligomeric intermediates are stable. **c,** Replication of the oligomeric intermediates is challenging. Replication 2 produced a similar spectral pattern to replication 1, indicating that MAS can induce oligomerization. The spectral pattern for replicate 3 was slightly different, indicating that it produced unstable oligomeric intermediates. The sample was 2 mM *h*IAPP in $D_2O$ (80 nmol *h*IAPP hydrated in 40 uL).

Unfortunately, when we attempted to reproduce the formation of the MAS-stabilized *h*IAPP oligomer, we struggled to replicate the data exactly (**Figure S2**). Defining the data in Figure 2 as Replicate 1, we performed the same experiment twice more. The Replicate 2 sample afforded $^{1}$H NMR lineshapes like those observed in Replicate 1 and exhibited similar monomer depletion kinetics. However, while Replicate 3 again displayed similar overall NMR lineshapes, the monomer depletion kinetics varied, showing ~40% depletion at 24 hours compared to 70-75% depletion in the first two trials. The peak near 0 ppm also reduced in intensity by ~40% in the third replicate, suggesting a degree of instability in the oligomer not seen in the first two replicates. No obvious explanation for the different behavior observed between the three replicates is immediately evident, but several possibilities exist. First, while the MAS frequency is maintained within 5-10 Hz (<0.1%) accuracy, there is less control over the rotor acceleration when MAS is initiated. Given that the oligomers seem to form in this initial spinning phase, it is possible that variations in the spin-up process may influence the range of *h*IAPP aggregates which immediately form. It is also possible that the *h*IAPP pretreatment with HFIP does not always perfectly dissolve preformed aggregates and that the formation of the MAS-induced oligomers is particularly sensitive to these preformed species. But we can only speculate based on the data reported here. Regardless, though the kinetics of monomer depletion and oligomer stability contrast slightly between trials, it is nevertheless reassuring that the nature of the oligomeric species does not appear to differ significantly, at least as can be seen in the $^1$H NMR lineshape.

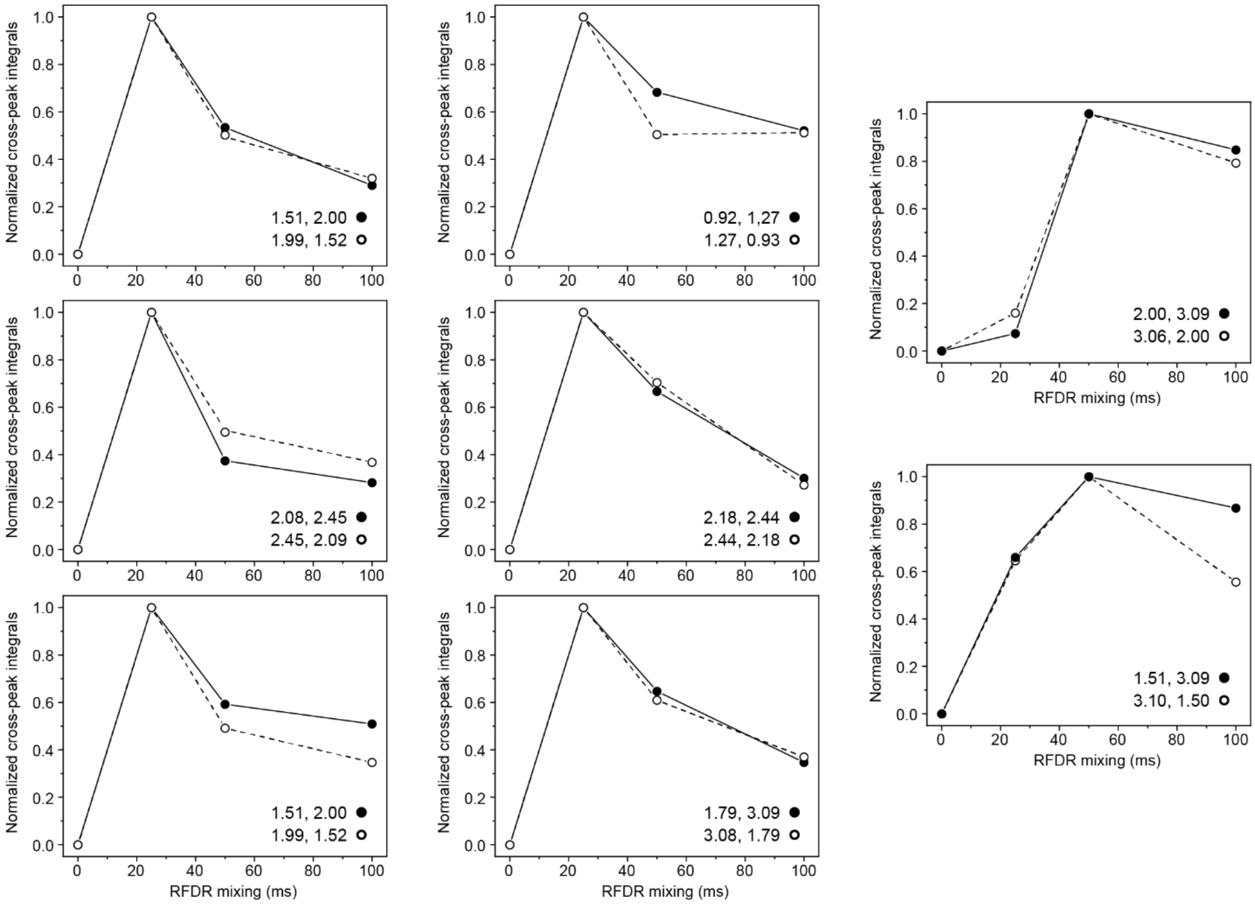


**Figure S3. RFDR Build-up curves for MAS-*h*IAPP in pure $D_2O$.** RFDR build-up curves are plotted for cross-peaks in the $^1$H-$^1$H NOE-RFDR spectra of 2 mM *h*IAPP in pure $D_2O$. NMR data was collected with an 800 MHz spectrometer and 15 kHz MAS.

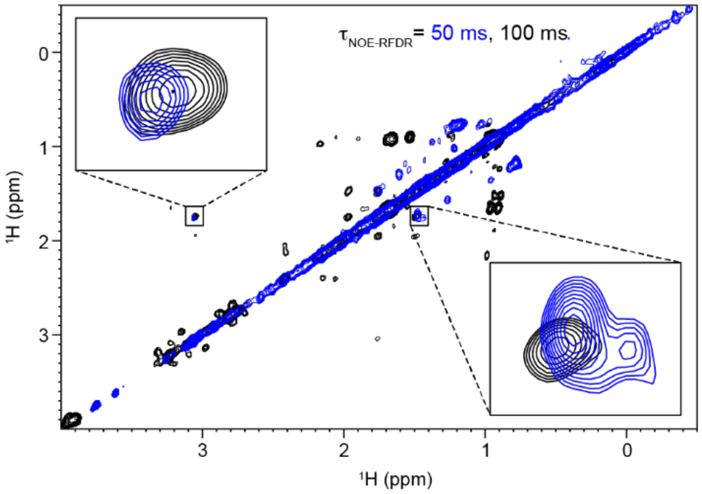


**Figure S4. Comparing dipolar recoupling from MAS-*h*IAPP in different solvent conditions.** 2D $^{1}$H-$^{1}$H NOE-RFDR spectra were obtained using a 500 MHz spectrometer with 10 kHz MAS at 298 K. The samples were 1 mM of *h*IAPP in 10 mM $d_3$-tris, 100 mM NaCl, pH 7.4 buffer (blue), or 1 mM *h*IAPP in
